# New Insights on the Evolutionary Relationships Between the Major Lineages of Amoebozoa

**DOI:** 10.1101/2022.02.28.482369

**Authors:** Yonas I. Tekle, Fang Wang, Fiona C. Wood, O. Roger Anderson, Alexey Smirnov

**Affiliations:** Dept. of Biology, Spelman College, Atlanta, Georgia, USA; Dept. of Biology and Paleo Environment, Lamont□Doherty Earth Observatory of Columbia University, Palisades, New York, USA; Dept. of Invertebrate Zoology, Faculty of Biology, St. Petersburg State University, Russia

**Keywords:** Amoebozoa, phylogenomics, flagellum, eukaryotes, genome, transcriptome

## Abstract

The supergroup Amoebozoa unites a wide diversity of amoeboid organisms and encompasses enigmatic lineages recalcitrant to modern phylogenetics. Deep divergences, taxonomic placement of some key taxa and character evolution in the group largely remain poorly elucidated or controversial. We surveyed available Amoebozoa genomes and transcriptomes to mine conserved putative single copy genes, which were used to enrich gene sampling and generate the largest supermatrix (824 genes) in the group to date. We recovered a well-resolved and supported tree of Amoebozoa, revealing novel deep level relationships and resolving placement of enigmatic lineages congruent with morphological data. In our analysis the deepest branching group is Tubulinea. A recent proposed major clade Tevosa, uniting Evosea and Tubulinea, is not supported. Based on the new phylogenetic tree, paleoecological and paleontological data as well as data on the biology of presently living amoebozoans, we hypothesize that the evolution of Amoebozoa probably was driven with the need to disrupt and graze on microbial mats - a dominant ecosystem of the mid-Proterozoic period of the Earth history.

## Introduction

The supergroup Amoebozoa ^1^ comprises a variety of amoeboid lineages; namely, naked lobose amoebae (which are “archetypal” amoebae), testate lobose amoebae, mycetozoans, anaerobic archamoebians and a heterogeneous assemblage of flattened amoeboid, branching reticulate or flagellated organisms; presently known as Variosea. Amoebozoa holds a key evolutionary position, being the closest known relative of Obazoa that, among other organisms, includes humans ^2,3^. Resolving the phylogenetic tree of this lineage is critical for answering important questions pertaining to the evolutionary origin of Amoebozoa, as well as for further clarification of the root of the eukaryotic tree ^3–8^.

Our understanding of the evolution and taxonomy of amoeboid protist originally conceived from cytological, morphological and life cycle evidence ^9,10^. Early studies based on small subunit rDNA (18S) gene indicated the polyphyly of naked amoebae (gymnamoebae) and formed the basis of our understanding of the supergroup Amoebozoa ^1,11,12^. The assemblage of Amoebozoa grew in membership, albeit with little improved resolution; or sometimes with conflicting hypotheses pertaining to within-group relationships (e.g., ^13–19^). This led to subsequent revisions and reevaluation in attempts to combine morphological and molecular characters and find synapomorphic characters of major clades ^20–23^. While this achieved major progress in our overall understanding of the group, much of the deep and intermediate relationships and placement of some groups of uncertain phylogenetic affinities (so-called *incertae sedis* taxa) remained elusive. Multigene studies, varying in breadth and depth of gene and taxon sampling, managed to overcome many of the challenges of single-gene reconstructions; and they resolved some of the long-standing evolutionary questions in the group ^4,24–27^. A recent phylogenomic study by Kang et al. ^4^ reported a deep level phylogeny of Amoebozoa based on large taxon sampling. However, the placements of some *incertae sedis* lineages were not entirely resolved. For some groups, other phylogenomic studies reported conflicting relationships ^25,26,28^.

The conflict in existing phylogenomic studies can be attributed partially to limitations of taxon and gene sampling as well as the methodology. Kang et al. ^4^ used large taxon sampling, but included only a small fraction of data (325 genes), from the vast amount of transcriptomic and genomic data available, based on commonly used genetic markers in eukaryotes. There are data suggesting that taxon sampling alone is not sufficient to resolve deep divergences in ancient lineages that might have undergone rapid radiations ^29^. The age of Amoebozoa is estimated to be over a billion years, and the probable origin of the group is dated back to the mid-Proterozoic period ^30,31^. Therefore, in order to infer deep evolutionary divergences not only increased taxon sampling, but also more representative genetic sampling along with the application of appropriate models and methods are essential.

In this study, we sampled putative single copy gene markers from genome-wide assays, increased taxon sampling and produced the largest amoebozoan supermatrix to date. This large dataset enabled us to recover a well-resolved and supported tree of the Amoebozoa. In addition, we uncover a well-corroborated novel deep-level relationship and resolved the placement of some *incertae sedis* lineages.

## Results

### The Tree of Amoebozoa

We recovered a monophyletic tree of Amoebozoa that is well resolved and supported in every one of our analyses (Figs. 1, S1-S4). Our datasets, with and without fast-evolving sites removed (analyzed using the complex model in IQ-TREE) recovered all well-established major subclades of Amoebozoa including Discosea, Archamoebae, Cutosea, Eumycetozoa, Variosea and Tubulinea with full support (Figs. 1, S1). The two well-known long-branch lineages, Archamoebae and Cutosea, were placed in their respective correct phylogenetic positions without removal of fast evolving sites in our full dataset (Fig. 1). Removal of fast evolving, rate categories, in IQ-TREE neither affected the topology nor improved support values (Fig. S1). In the RAxML analysis, the accurate placement of Archamoebae and Cutosea, required removal of six fast evolving rate categories (38%) from the full dataset (Fig. S2); but resulted in the same final tree configuration. The RAxML tree had generally lower supported branches but was congruent with the topology of the trees inferred using IQ-TREE (Figs. 1, S1, S2). A similar reduced dataset was analyzed using Bayesian inference, which yielded similar topology despite lack of convergence in our PhyloBayes analysis (data not shown). Kang et al. ^4^ also reported similar topologies among their ML and PhyloBayes trees despite limited number of chains used and lack of convergence in some of their PhyloBayes analyses. Due to the high computational demand, Bayesian inference was not feasible with our large dataset. The consistency of tree topologies across methods and algorithms used, as well as the placement of long-branch taxa (Archamoebae and Cutosea) without removal of fast evolving sites in IQ-TREE (likely due to complex model used), demonstrates the robustness of our result.

**Figure 1.**
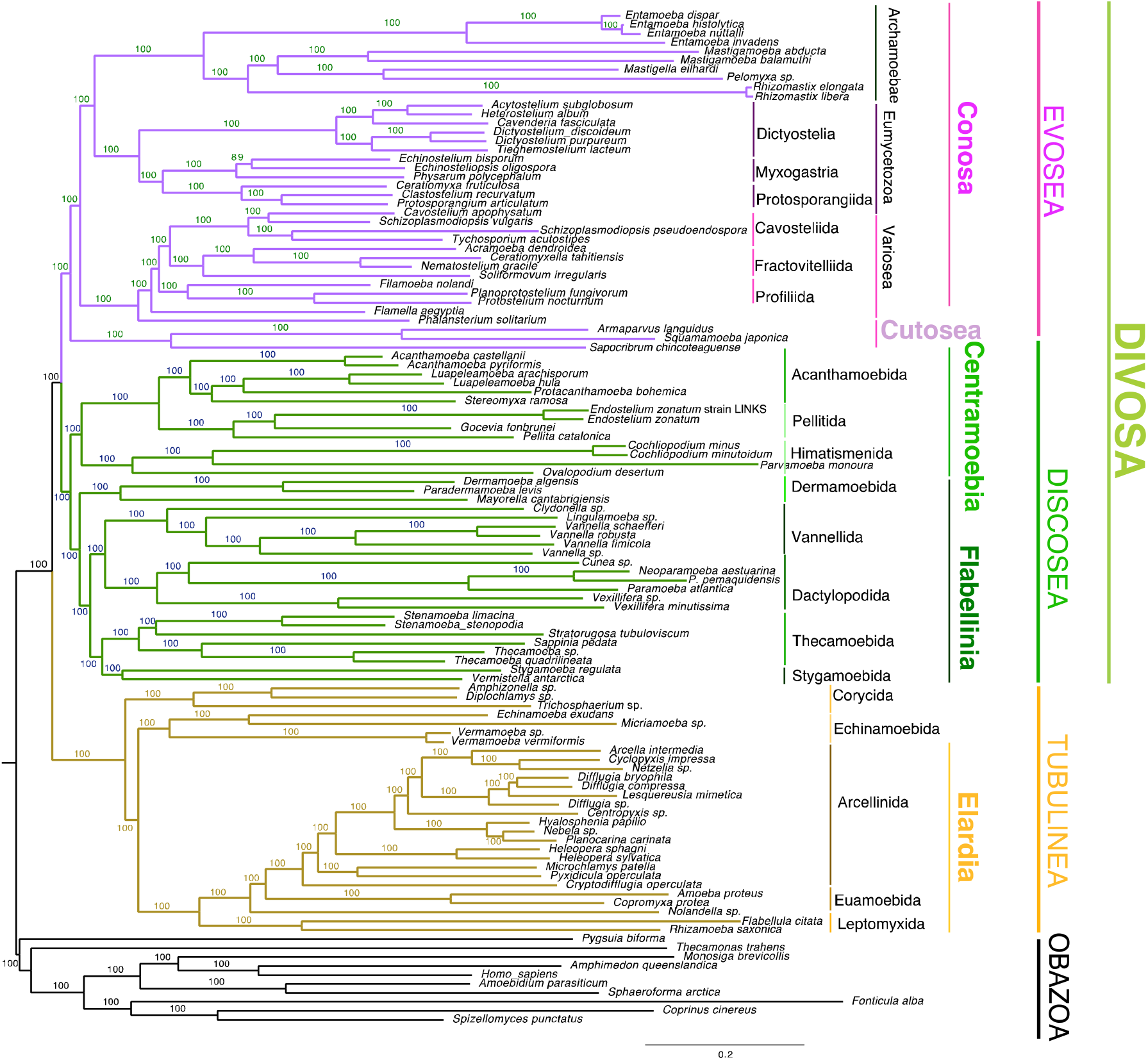
Genome wide phylogeny of the Amoebozoa inferred using Maximum likelihood (ML) in IQ-TREE with LG+G4+C60+F model of evolution. The data matrix used to infer this tree consisted of 113,910 amino acid sites from the full dataset, derived from 824 genes and 113 taxa including 10 outgroup taxa. Clade supports at nodes are ML IQ-TREE 1000 ultrafast bootstrap values obtained using the same model. All branches are drawn to scale except a branch leading to Archamoebae, and *Sapocribrum chincoteaguense* and *Parvamoeba monoura*, that were reduced to one-third and half, respectively.

In our phylogenomic tree, all major clades are congruent with previous published topologies ^4,24–26^. Moreover, our phylogenomic tree has well-corroborated relationships; and the recovery and placement of enigmatic taxa are more stable (Figs. 1, S1, S2). Our results yielded improved support for the Flabellinia and Thecamoebida clades compared to a previous comparable phylogenomic study ^4^. We have recovered for the first time a fully supported monophyletic clade encompassing two *incertae sedis* taxa, *Vermistella* and *Stygamoeba*. Both these lineages were placed in the order Stygamoebida based on morphological evidence ^22^. The monophyly and placement of this order in the tree of Amoebozoa has not been resolved in previous multigene analyses (e.g., ^4^). In our tree Stygamoebida clade forms a sister group relationship with Thecamoebida with full support (Fig. 1). We also find some discrepancies between our tree (Fig. 1) and that of Lahr et al. ^5^ in the branching order of the Tubulinea clade, albeit with similar taxon sampling for this clade. Our analysis shows clade Corycida as the most basal Tubulinea lineage similar to that of Kang et al. ^4^ phylogeny (Fig. 1). *Nolandella* sp., a member of Euamoebida, did not group with *Amoeba proteus* and *Copromyxa protea* in our analysis, but formed an independent lineage (Fig. 1).

### A Novel Deep Split of the Amoebozoa

Our analysis for the first time revealed a novel, well-supported deep spilt of Amoebozoa; not reported in previous phylogenomic studies. Amoebozoa is split into two fully supported major subclades: Tubulinea and a second one comprised of the remaining major subclades including Evosea (Eumycetozoa, Variosea, Archamoebae, and Cutosea) and Discosea (Figs. 1, S1, S2). This branching is different from a finding in a recent phylogenomic study that reported a spilt between Discosea and Tevosa (Evosea+ Tubulinea) ^4^. Tevosa is not supported in our analyses, including analyses with removal of fast sites. On the other hand, the deep split (Evosea+Discosea *vs*. Tubulinea) observed in our phylogenomic tree is supported in all analyses of our data sets. The deep spilt receives almost full support in our internode certainty (IC) analyses as implemented in QuartetScores (1.00) and RAxML (0.979) (Figs. S3, S4). AU test of our topology, comparing alternative topologies with Tevosa and a traditional deep relationship uniting Discosea and Tubulinea (Lobosa), showed that the newly recovered deep spilt has the highest p-value (p-AU = 0.947). Hypothesis Lobosa was rejected (p-AU = 0.000278), while Tevosa cannot be rejected with p-value just above threshold (p-AU = 0.0564). For convenience, we suggest a new name for the deep spilt (Discosea+Evosea) clade; i.e., Divosa, a term derived from a combination of the name of the two clades.

## Discussion

### Targeted Genome-Wide Data Enrichment for Phylogenomics of Amoebozoa

Despite the large number of RNA-Seq data generated in recent studies ^4,24–26^, only a small fraction of this data has been utilized in phylogenomic analyses. To increase it, we compiled a total of 1559 markers using genome-derived protein coding genes from 113 amoebozoan genomes and transcriptomes. Using putative single copy markers, primarily derived from Amoebozoa genomes, has enabled us to introduce highly conserved markers with phylogenetic signal corroborating morphology- and phylogenomic-based amoebozoan hypotheses ^4,24^. While single-copy genes identified in some genomes might not always apply to others, a previous phylogenomic study with seed plants, based on single copy markers resulted in more resolved phylogeny both at shallow and deep nodes ^32^. In this study, we followed a stringent approach aided by automated and manual curation of markers, selected from the above-mentioned dataset to build the largest supermatrix (823 genes) in the Amoebozoa. With this approach, we substantially increased the total number of genes used in Amoebozoa phylogenomics. Our analysis yielded consistent and well-corroborated topologies, despite whether we included or excluded fast evolving sites (Figs. 1, S2). The robustness of our phylogeny is also corroborated with the high support values from internode certainty analysis (Figs. S3, S4). One of the evident results of this approach is the first time phylogenomic recovery of the monophyly of the taxon Stygamoebida, earlier supported only at the morphological level ^22,23^ and a recovery of a novel deep split divergence of Amoebozoa.

### Unraveling deep divergence of Amoebozoa

A recent phylogenomic study by Kang et al. ^4^, though based on a slightly smaller taxon sampling, proposed a split of the Amoebozoa supergroup into two major subclades: Tevosa (Evosea+Tubulinea) and Discosea. By contrast, in our study Evosea robustly groups as sister clade to Discosea (Figs. 1, S1, S2). Both phylogenetic hypotheses, ‘Tevosa’ and Divosa, receive high statistical support in their and our study, respectively (see Fig. 1, ^4^). In phylogenomic analyses, it is common to see that short subtending deep nodes receive high statistical support ^33^. Amoebozoan deep nodes are characterized by very short branch lengths, an indication of limited supporting characters, or possible ancient rapid diversification. Strong statistical support at these levels of nodes does not necessarily mean that the inferred relationships are correct. Statistical indices such as bootstrap values and Bayesian posterior probabilities only assess sampling effects, and give an indication of tree reliability that is dependent on the data and the method ^34^. This can partially explain why these short-branch, deep nodes in Amoebozoa phylogenomic studies tend to collapse, or vary, depending on the method of analysis or the composition of the gene/taxon sampling ^4,24–26^. Certainly, caution still must be taken when interpreting ancient divergences, because results can be muddied by noise (e.g., gene history ^35^ or lack of signal due to rapid radiation ^29^). However, the support of the split recovered in the present study is high and originates from different lines of evidence.

It is possible to note that in many lineages trophozoites of Discosea and Variosea are more similar to each other rather than to Tubulinea. Certainly, the morphology of presently living amoeboid organisms is derived and adaptive, but generally it is possible to say that members of Divosa lineage share more morphological similarity between each other rather than with the Tubulinea lineage. For example, amoebae of the genus *Flamella*, belonging to the class Variosea, by their morphology may be easily confused with some discosean amoebae (e.g., ^36^); the same is true for individual trophozoites of many mycetozoan species, showing flattened body shape and pointed subpseudopodia ^37,38^. Cells of amoebae belonging to the genus *Squamamoeba* (the taxon of Cutosea), sometimes resemble *Korotnevella* (Discosea) in their overall morphology, hence, being differently organized at the cytological level ^39^. At the same time, none of discosean or variosean lineages show the morphology resembling that of, e.g. Amoebida, or alteration of the locomotive morphology from flattened to tubular, which is a general characteristic of Tubulinea ^20,22^. To certain extent, the return to the tubular body shape, subcylindrical in cross-section occurs among amoeboid representatives of Archamoebea; however, this might be mostly related with their specific lifestyle (parasites or pelobionts). In addition the pattern of pseudopod formation (e.g., the tendency to show eruption of the hyaline cytoplasm in the frontal area of the cell) makes them to be significantly different from that in Tubulinea (see ^40^).

### Mid-Proterozoic environment – the driving force for the origin of Amoebozoa

The flagellum (cilium) is a highly conserved complex structure that is believed to have originated only once, and be ancestral to all eukaryotes ^2,41,42^. Amoebozoa are remarkable in that the two basal phylogenetic lineages, Tubulinea and Discosea, have entirely lost cilia, kinetosomes (basal bodies) and associated root structures; while a derived major clade, Evosea, contains a handful of ciliated lineages in a few branches intermingled among amoeboid lineages ^21,22^. The loss of cilia and associated structures in the majority members of Amoebozoa is one of the biggest mysteries pertaining to their origin and evolution.

In ciliated members of Amoebozoa, the ciliary apparatus is characterized by a specific arrangement of root structures, which includes an incomplete (Variosea and Mycetozoa) or complete (Archamoebea) cone of microtubules extending from the kinetosome to the nucleus ^43^. In early interpretations, this conical arrangement of microtubules was considered to be homologous to the ciliary root system of Opisthokonta; which, together with other morphological and molecular evidence, gave rise to the “Unikonta” hypothesis ^2,44,45^. In this model, the hypothetical ancestor of Amoebozoa was considered to be an organism with a single emergent cilium, resembling *Phalansterium* or *Mastigamoeba* in cellular organization ^46,47^. This lineage, combining Amoebozoa and Opisthokonta, has been proposed as an alternative to that of the bikonts, with two emerging cilia; which included the rest of the eukaryotic groups. Cavalier-Smith ^2^ argued that among unikonts, paired kinetosomes (when present) resulted from convergent evolution rather than common ancestry with bikonts. Molecular and morphological analyses provided certain indications that the microtubular structures in Amoebozoa, and Opisthokonta may not be homologues ^43,48^. However, further development of molecular phylogeny provided evidence for the basal position of bikont organisms in the tree of eukaryotes ^3,49,50^. Thereafter, the general consensus nowadays is that hypothetical common ancestor of Amoebozoa, was a bikont organism ^43,51,52^. Several authors (e.g., ^3,43,49,50^) hypothesised that the presumable common ancestor was a ventrally grooved biciliate gliding flagellate, capable of producing filose ventral pseudopodia and possessing a relatively complex organization of the cell. That is, a cell possessing two cilia with kinetosomes and root structures, ventral groove supported with microtubules and dorsal pellicle – the so called “sulcozoan ancestor”. Its name originates from Sulcozoa – a phylum of protists established by Cavalier-Smith ^43^ that combines a heterogenous assemblage of early evolving eukaryotic lineages. Cavalier-Smith suggested that “opisthokonts and Amoebozoa evolved from sulcozoan ancestors by two independent losses of the pellicular dense layers and of the ventral groove, which in both cases would allow pseudopods to develop anywhere on the cell surface” (op. cit.).

The origin and further evolution of Amoebozoa in this hypothesis presumes the loss of both cilia and kinetosomes in Lobosa (Tubulinea and Discosea) and of the posterior cilium and one kinetosome in most of the ancestors of Conosa - Archamoebae, Variosea and Eumycetozoa; Cutosea were not known at that time (e.g., ^3,49,50^). This evolutionary scenario was rather logical and is illustrated in Figure 2A. However, the Lobosa/Conosa dichotomy was doubted based on some 18S gene phylogenies ^27^; and it subsequently failed to garner support in wide-scale phylogenomic studies ^4,24,25^, as well as in the present study. This makes the model of multiple losses more complicated, because under the new tree configuration, we have to suggest subsequent partial or complete loss of cilia and related structures in all but one branch of Amoebozoa. This hypothetical scenario is illustrated in Figure 2B. It remains unclear why the hypothesized ancestor of Amoebozoa, being initially a quite complex biciliated organism, underwent such a massive loss (or substantial simplification) of cilia-related structures in almost all evolutionary lineages of Amoebozoa, and what was the driving force for such a reduction.

**Figure 2.**
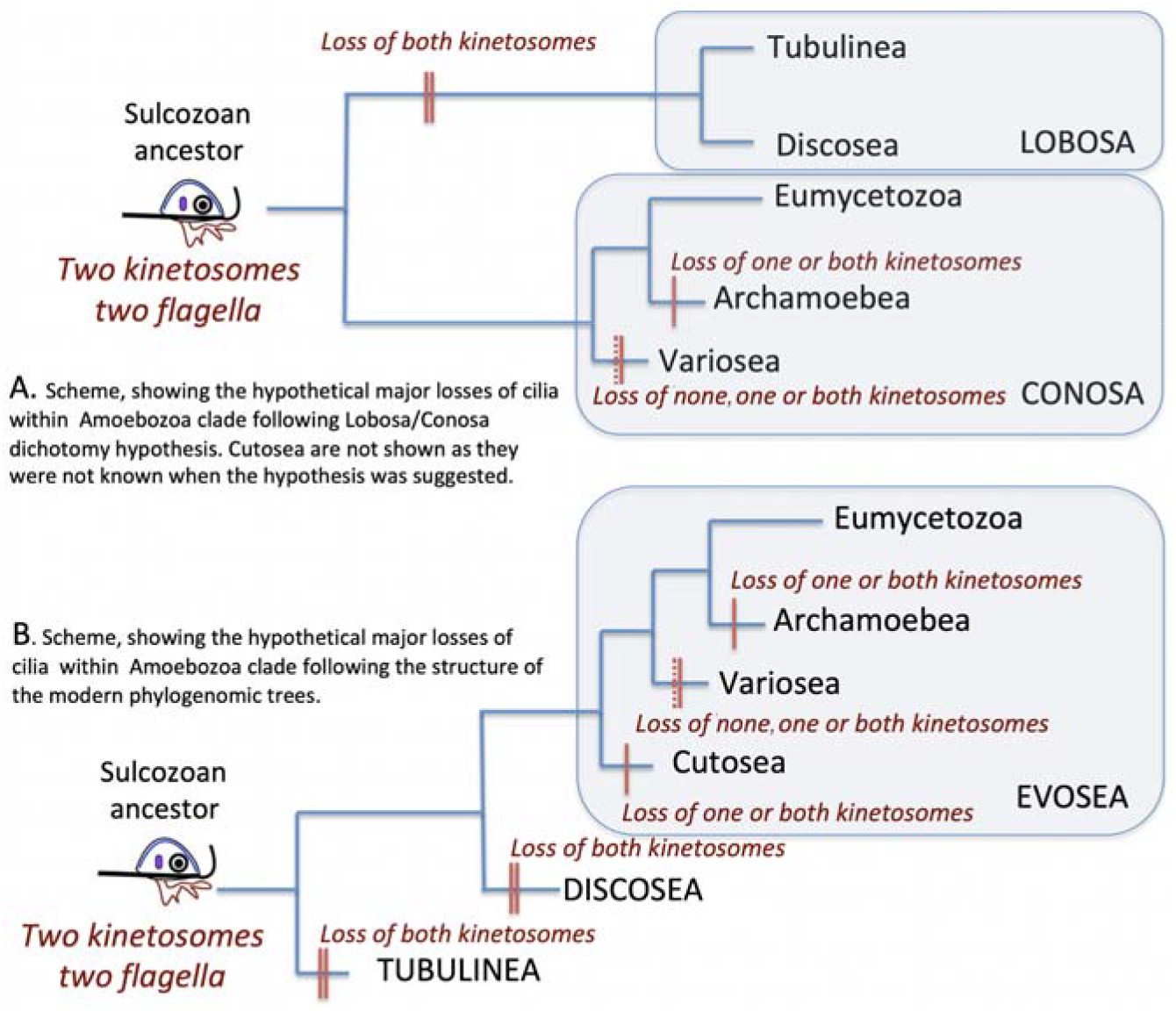
A scheme illustrating the loss of kinetosomes and cilia under the different evolutionary hypotheses (A and B). Vertical hash marks on branches show loss of kinetosomes (the number lost as designated by labels on the diagram) depending on the lineage.

Several studies based on molecular dating analysis correspondingly placed the origin of Amoebozoa to the Mesoproterozoic period, which means 1250 – 1624 mya ^31,53^. It means that the early evolution of Amoebozoa took place at the period when the biosphere was dominated with microbial biofilms – sheets of bacteria, embedded in extracellular polymeric substances, covering almost every possible substrate ^54^. Being initially rather simple, biofilms further evolved in complex microbial mats, comprising different prokaryotic organisms, showing concerted activities and intimate interactions between various microbial metabolisms ^55^. The oldest mats are dated to approximately 3.5 billion years ago, and the noonday of mats covers the mid-Proterozoic period ^56,57^, which roughly corresponds to the estimate of the potential age of Amoebozoa.

Formation of a microbial biofilm, among other structural and biogeochemical features, can be explained as an adaptation that increases survival of bacteria to avoid predation ^58,59^. The probable size of the bacterivorous biflagellate ancestor of Amoebozoa was relatively small, likely no larger than that of the existing representatives of the CRuMs clade (e.g., *Mantamonas*) or ‘Excavates’ (metamonads or *Malawimonas*), which is within the general size range of 2-20 μm. These organisms were able to phagocytize solitary bacteria, but consumption of microorganisms embedded in an intact microbial mat probably was beyond their capacity, as well as this is beyond the capacity of the modern flagellates of comparable size ^60,61^. Feeding on bacteria, major constituents of the microbial mats (the dominant food source in the mid-Proterozoic environment), required increment in the body size and acquisition of special adaptations allowing them to ingest filamentous food. However, the latter was again related to the body size, because the filament, even compacted in some way, must be ingested – i.e., appear inside the cell.

Due to Reynolds number limitation ^62,63^, the increment in the body size makes ciliary motility less adaptive due to loss of efficiency. Thus, from an adaptive aspect, an amoeboid lifestyle might be a way to increase the body size while retaining a motility function, no longer dependent on cilia. An amoeboid organization also could gain the adaptive capacity to disrupt microbial mats and graze, feeding on bacteria within the mats. This adaptation would provide access to the dominant food source in the biosphere of the mid-proterozoic eon. Indeed, presently, naked amoebae are known as one of the primary grazers of bacterial biofilms ^64–66^. Moreover, they not only just graze and phagocytize prey in the mats, but also disrupt them, making their content available for other organisms ^67,68^. Finally, in addition to the advantage of feeding on bacterial mats ^69,70^, it is also possible that an increase in body size alleviated pressure of predation by other organisms on the last Amoebozoan common ancestor (LACA), which for some time provided it an adaptive advantage and allowed rapid proliferation and differentiation of Amoebozoa in the mid-Proterozoic environment.

Hence, we hypothesise that the adaptive value of amoeboid locomotion and concomitant grazing potential on the dominant food source in the mid-proterozoic biosphere – the microbial mats – favoured the evolution of the Amoebozoa. They probably successfully solved this task by the increment of body size. However, at the same time, the efficiency of flagellar locomotion was highly reduced or lost; and this resulted in the multiple suspensions of the flagellar apparatus, which is completely absent in two major current amoebozoan lineages – Tubulinea and Discosea (Fig. 2). The modern configuration of the Amoebozoan tree, which rejects the Lobosa/Conosa dichotomy and suggests a subsequent branching of lineages (with either Tubulinea or Discosea at the base), leaves open a major question. That is, was the last Amoebozoa common ancestor an amoeboflagellate, with the domination of amoeboid movement based on the microtubular cytoskeleton; or was the flagellum-related structures and microtubular locomotive system entirely suppressed? If the latter case is true, then it probably drove the ancestral amoebozoan to switch to the acto-myosin movement, as found in modern representatives of naked and testate lobose amoebae. Probably, the answer to this question may be obtained by the analysis of gene content and the level of flagellum-related gene expression in the amoebozoan genomes. However, the dataset available for quality analysis remains limited in this group of protists and requires further accumulation prior to conclusive study.

## Methods

### Transcriptome Assembly and Contamination Examination

All transcriptome data used in this study were assembled using a bioinformatics pipeline described in Tekle and Wood ^25^. As a precautionary measure for contamination, high-quality data generated from single cell or monoclonal cultures, and without history of contamination, were prioritized in our data collection. We also checked highly conserved genes (e.g., small subunit rDNA and cytoskeletal genes) for assembled transcriptomes to check the identity of the species. Species suspected to have been contaminated (e.g., *Ripella* sp. DP13-Kostka) or with low- or poor-quality transcriptome data (see below) have been removed from the final analysis. Assembled contigs were translated into protein sequences using TransDecoder (https://github.com/TransDecoder/TransDecoder/wiki).

### Taxon and gene sampling

A total of 107 amoebozoans representing the vast diversity of the supergroup and 10 outgroup taxa from a closely related clade, Obazoa, were included in our initial analysis (Table S1). Four ingroup taxa including *Parvamoeba rugata*, *Centropyxis aculeata*, *Hyalosphenia elegans* and *Grellamoeba robusta*, were removed from the final dataset due to poor data quality. A recent phylogenomic study ^5^ that focused on testate amoebae (clade Tubulinea) reported a topology of Tubulinea that differed from that of Kang et al. ^4^. To explore these discrepancies further, and assess the impact of taxon sampling on branching order of Tubulinea clade and its position within the Amoebozoa phylogeny, we added more slowly evolving taxa to Tubulinea. The final supermatrix consisted of 113 taxa including the outgroup taxa (Table S1).

A genome wide gene sampling approach using available amoebozoan genomes was employed to identify single copy markers. Previous phylogenomic studies have used conserved phylogenetic markers commonly found in a wide range of eukaryotic diversity ^4,24^. In this study we used a series of bioinformatics steps to maximize gene sampling in the Amoebozoa. We conducted a whole genome comparison of three well-annotated amoebozoan genomes, *Acanthamoeba castellanii*, *Dictyostelium discoideum* and *Entamoeba histolytica*, to extract commonly shared protein-coding genes among these genomes in OrthoVenn ^71^. Inclusion of *E. histolytica* greatly reduced the number of shared genes by 40% because this amitochondriate parasitic species has a comparably much reduced genome to the free-living amoebae. For this reason, to be more representative, further comparative analysis was done using *A. castellanii* and *D. discoideum* as reference genomes to mine single-copy genes. Using this approach, we identified 1559 putative single copy genes that were used as a query to search orthologous genes from ingroup and outgroup taxa.

We used NCBI-BLAST with e-value threshold of 10^-15^ to retrieve homologous sequences from transcriptomes or genomes of our selected taxa. From this analysis, sequences with best e-value scores were retained for each taxon. The retained sequences, for each taxon and gene, were compiled and aligned using a sequence alignment tool, MAFFT, with default setting ^72^. These alignments were then trimmed in TrimAl ^73^ using “automated1” setting provided by the program. To inspect potential paralogs from each gene, we inferred single gene trees using IQ-TREE with the best-fit model automatically fast selected by ModelFinder ^74^. Both single gene trees and their corresponding alignments were then inspected manually for paralogy and other anomalies related to alignment accuracy, sequence length and fast evolving lineages ((Single gene alignment and trees available for review on this link: https://www.dropbox.com/). We applied strict gene selection criteria that included removal of anomalous grouping (e.g., lineages that grouped with outgroup or wrong (unexpected) phylogenetic position with >90% bootstrap support) and genes that showed paralogy (duplication) signs. To mitigate the impact of long-branch attraction during phylogenetic reconstruction, we removed genes that contained three or more long-branch lineages. Two exceptions for this approach were the well-known long-branch lineages, Cutosea and *Entamoeba*, that were kept in all of our analyses. These two lineages were retained since all their representatives are mostly long-branches. They are also indirect indicators of noise in a data matrix since their correct placement usually requires removal of fast-evolving sites due to the effect of long-branch attraction. Following these criteria, we retained a total of 824 gene clusters in the final dataset. Orthologous group numbers were assigned for each gene cluster using ublast in USEARCH ^75^ with e-value 10^-10^. We used the OrthoMCL database to generated ortholog group numbers ^76^ (Table S2).

### Supermatrix Construction and Tree Inference

The alignments from 824 genes were concatenated into an initial supermatrix containing 198,280 amino acid sites and 117 taxa using a customized R script. Taxa with over 80% gappy sites were removed, which resulted in exclusion of 4 lineages (*Parvamoeba rugata*, *Centropyxis aculeata*, *Hyalosphenia elegans*, *Grellamoeba robusta*). Constant sites, and sites with more than 50% missing data, were removed from this alignment, and the resulting supermatrix retained 113,910 amino acid sites and 113 taxa for the full dataset.

Phylogenomic analyses of the final datasets were conducted in IQ-TREE – an ^74^ efficient tool to analyze large datasets by the maximum likelihood (ML) method. All IQ-TREE analyses were preformed using LG+G4+C60+F model, with 1000 replicates for ultrafast bootstrap, which allowed full profile mixture model C60 and Gamma rate heterogeneity across sites. We also analyzed our dataset in RAxML v.8.2.X ^77^ using PROTGAMMALG4X model; branch support was estimated from 1000 rapid bootstrap pseudoreplicates.

Fast-evolving sites and taxa are known to be problematic for tree inference due to saturation of substitutions and subsequent convergent evolution resulting in long-branch attraction (LBA) and other systematic errors. To test the effects of these types of errors on our phylogenomic analysis, we performed a site removal assay in which each site of the supermatrix was assigned to one of 16 categories based on its rate from IQ-TREE. This was performed using a posterior mean site frequency (PMSF) model with mixture model C60 and 16 discrete rate categories of sites. For this analysis, we used the tree from full dataset inferred above as a guide tree. The impact of fast evolving sites on resulting phylogenies was assessed by subsequent removal of fast categories of sites (up to 6 categories). In IQ-TREE our full dataset was analyzed with 3 categories removed using PMSF model with a guide tree inferred from the complex model (LG+G4+C60+F) mentioned above. In RAxML, 3 and 6 fast site categories were removed and analyzed using the same model as above.

### Internode Certainty Analysis and Hypothesis Testing

As alternative to bootstrap branch support from IQ-TREE, we calculated internode certainty (IC) scores using the program QuartetScores ^78^. This approach calculated IC scores from the frequencies of quartets, which can correct for the missing taxa using a set of trees. For this analysis, we used 1000 bootstrap trees generated from LG+G4+C60+F model in IQ-TREE with our full dataset. Alternatively, we used RAxML to estimate the degree of certainty for internodes and tree topology for bipartitions with PROTGAMMALG4X model ^79^.

We used Approximately Unbiased (AU) tests ^80^ to test alternate tree topologies pertaining to the deep node hypotheses Divosa (this study), Tevosa (Kang et al. 2017) and Lobosa ^27^ with the full dataset (113,910 sites). Two loosely constrained topologies Tevosa ([Tubulinea+Evosea]+Discosea) and Lobosa ([Discosea+Tubulinea]+Evosea) were optimized under LG+G4+F+C60 in IQ-TREE. These optimized trees were compared with our tree (Divosa, ([Discosea+Evosea], Tubulinea) using AU test with 10,000 RELL bootstrap replicates ^81^. The hypotheses that had p-AU ≥ 0.05 within the 95% confidence interval could not be rejected.

## Acknowledgments

This work is supported by the National Science Foundation EiR (1831958) and National Institutes of Health (1R15GM116103-02) to YIT. Additional support was gained from RSF 20-14-00195 to AS (evolutionary analysis). We would like to thank James T. Melton III, Estifanos Zerai, Ludmila Chystyakova, Sergei Karpov and Mandakini Singla for assistance in data collection, preliminary analysis and general discussions.

## Author contributions

YIT conceived the project, led writing manuscript and helped design experiments and analysis. FW and FCW collected data, conducted analysis, and contributed to writing and editing of the manuscript. ORA and AS helped with writing, editing and organizing of the manuscript. All authors have read and approved the manuscript.

## Supplementary Figure caption

**Figure S1**. Genome wide phylogeny of the Amoebozoa inferred using Maximum likelihood (ML) in IQ-TREE with LG+G4+C60+F model of evolution. The data matrix used to infer this tree consisted of 93,820 sites amino acid sites with three fast categories of sites (13%) removed from the full dataset. The data matrix consists of 824 genes and 113 taxa including 10 outgroup taxa. The topology was estimated under LG+G4+C60+F+PMSF [Y1] model using a guide tree from a topology estimated using full dataset shown in Figure 1. Clade supports at nodes are ML IQ-TREE 1000 ultrafast bootstrap values obtained using the same model. All branches are drawn to scale.

**Figure S2**. Maximum Likelihood tree inferred by RAxML with six fast categories of sites removed from the full dataset. The topology was estimated under PROTGAMMALG4X model. Total number of sites included after removing six fast sites categories is 70,543.

**Figure S3**. Internode certainty inferred by QuartetScores for topology in Figure 1. Values at branches are Quadripartition internode certainty (qp-ic); Lowest quartet internode certainty (lp-ic); Extended Quadripartition internode certainty (eqp-ic).

**Figure S4**. Internode certainty inferred using RAxML under PROTGAMMALG4X model for topology in Figure 1. Branch labels showed the internode certainty for a given internode with the most conflicting bipartition (left value) or all conflicting bipartitions (right value). Relative tree certainty including all conflicting bipartitions for this tree is 0.978410.

## References

1 Cavalier-Smith, T. A revised six-kingdom system of life. Biological Reviews of the Cambridge Philosophical Society 73, 203–266 (1998).

2 Cavalier-Smith, T. The phagotrophic origin of eukaryotes and phylogenetic classification of protozoa. International Journal of Systematic and Evolutionary Microbiology 52, 297–354 (2002).

3 Brown, M. W. et al. Phylogenomics demonstrates that breviate flagellates are related to opisthokonts and apusomonads. Proc Biol Sci 280, 20131755, doi:10.1098/rspb.2013.1755 (2013).

4 Kang, S. et al. Between a Pod and a Hard Test: The Deep Evolution of Amoebae. Mol Biol Evol 34, 2258–2270, doi:10.1093/molbev/msx162 (2017).

5 Lahr, D. J. G. et al. Phylogenomics and Morphological Reconstruction of Arcellinida Testate Amoebae Highlight Diversity of Microbial Eukaryotes in the Neoproterozoic. Curr Biol 29, 991–1001 e1003, doi:10.1016/j.cub.2019.01.078 (2019).

6 Yoon, H. S. et al. Broadly sampled multigene trees of eukaryotes. BMC Evolutionary Biology 8, 14 (2008).

7 Burki, F. et al. Phylogenomics reshuffles the eukaryotic supergroups. PLoS ONE 2, e790 (2007).

8 Parfrey, L. W. et al. Broadly sampled multigene analyses yield a well-resolved eukaryotic tree of life. Systematic biology 59, 518–533, doi:syq037 [pii] 10.1093/sysbio/syq037 (2010).

9 Page, F. C. The classification of ‘naked’ amoebae (Phylum Rhizopoda). Arch. Protistenkd. 133, 199–217 (1987).

10 Rogerson, A. & Patterson, D. J. The Naked Ramicristate Amoebae (Gymnamoebae). In: Lee, J.J., Leedale, G.F., Bradbury, P. (Eds.), An Illustrated Guide to the Protozoa, 2nd ed. Society of Protozoologists, Lawrence, Kansas, pp. 1023–1053 (2002).

11 Amaral Zettler, L. A. et al. Microbiology: Eukaryotic diversity in Spain’s River of Fire. Nature 417, 137 (2002).

12 Cavalier-Smith, T. & Chao, E. E. Molecular phylogeny of the free-living archezoan Trepomonas agilis and the nature of the first eukaryote. Journal Of Molecular Evolution 43, 551–562 (1996).

13 Tekle, Y. I. et al. Phylogenetic placement of diverse amoebae inferred from multigene analyses and assessment of clade stability within ‘Amoebozoa’ upon removal of varying rate classes of SSU-rDNA. Molecular phylogenetics and evolution 47, 339–352 (2008).

14 Fahrni, J. F. et al. Phylogeny of lobose amoebae based on actin and small-subunit ribosomal RNA genes. Mol Biol Evol 20, 1881–1886, doi:10.1093/molbev/msg201 (2003).

15 Nikolaev, S. I. et al. The testate lobose amoebae (order Arcellinida Kent, 1880) finally find their home within Amoebozoa. Protist 156, 191–202 (2005).

16 Fiore-Donno, A. M., Meyer, M., Baldauf, S. L. & Pawlowski, J. Evolution of dark-spored Myxomycetes (slime-molds): molecules versus morphology. Mol Phylogenet Evol 46, 878–889, doi:10.1016/j.ympev.2007.12.011 (2008).

17 Berney, C. et al. Expansion of the ‘Reticulosphere’: Diversity of Novel Branching and Network-forming Amoebae Helps to Define Variosea (Amoebozoa). Protist 166, 271–295, doi:10.1016/j.protis.2015.04.001 (2015).

18 Amaral Zettler, L. A. et al. A molecular reassessment of the leptomyxid amoebae. Protist 151, 275–282 (2000).

19 Bolivar, I., Fahrni, J. F., Smirnov, A. & Pawlowski, J. SSU rRNA-based phylogenetic position of the genera *Amoeba* and *Chaos* (Lobosea, Gymnamoebia): The origin of gymnamoebae revisited. Molecular Biology and Evolution 18, 2306–2314 (2001).

20 Smirnov, A. et al. Molecular phylogeny and classification of the lobose amoebae. Protist 156, 129–142 (2005).

21 Cavalier-Smith, T., Chao, E. E. Y. & Oates, B. Molecular phylogeny of Amoebozoa and the evolutionary significance of the unikont *Phalansterium*. European Journal of Protistology 40, 21–48 (2004).

22 Smirnov, A. V., Chao, E., Nassonova, E. S. & Cavalier-Smith, T. A revised classification of naked lobose amoebae (Amoebozoa: lobosa). Protist 162, 545–570, doi:S1434-4610(11)00031-9 [pii]10.1016/j.protis.2011.04.004 (2011).

23 Adl, S. M. et al. Revisions to the Classification, Nomenclature, and Diversity of Eukaryotes. J Eukaryot Microbiol 66, 4–119, doi:10.1111/jeu.12691 (2019).

24 Tekle, Y. I. et al. Phylogenomics of ‘Discosea’: A new molecular phylogenetic perspective on Amoebozoa with flat body forms. Mol Phylogenet Evol 99, 144–154, doi:10.1016/j.ympev.2016.03.029 (2016).

25 Tekle, Y. I. & Wood, F. C. Longamoebia is not monophyletic: Phylogenomic and cytoskeleton analyses provide novel and well-resolved relationships of amoebozoan subclades. Mol Phylogenet Evol 114, 249–260, doi:10.1016/j.ympev.2017.06.019 (2017).

26 Cavalier-Smith, T., Chao, E. E. & Lewis, R. 187-gene phylogeny of protozoan phylum Amoebozoa reveals a new class (Cutosea) of deep-branching, ultrastructurally unique, enveloped marine Lobosa and clarifies amoeba evolution. Mol Phylogenet Evol 99, 275–296, doi:10.1016/j.ympev.2016.03.023 (2016).

27 Cavalier-Smith, T. et al. Multigene phylogeny resolves deep branching of Amoebozoa. Molecular phylogenetics and evolution 83, 293–304, doi:10.1016/j.ympev.2014.08.011 (2015).

28 Tekle, Y. I. & Williams, J. R. Cytoskeletal architecture and its evolutionary significance in amoeboid eukaryotes and their mode of locomotion. R Soc Open Sci 3, 160283, doi:10.1098/rsos.160283 (2016).

29 Shin, S. et al. Taxon sampling to address an ancient rapid radiation: a supermatrix phylogeny of early brachyceran flies (Diptera): Diptera evolution and supermatrix. Systematic Entomology, doi:DOI: 10.1111/syen.12275 (2017).

30 Eme, L., Sharpe, S. C., Brown, M. W. & Roger, A. J. On the age of eukaryotes: evaluating evidence from fossils and molecular clocks. Cold Spring Harb Perspect Biol 6, doi:10.1101/cshperspect.a016139 (2014).

31 Parfrey, L. W., Lahr, D. J., Knoll, A. H. & Katz, L. A. Estimating the timing of early eukaryotic diversification with multigene molecular clocks. Proc Natl Acad Sci U S A 108, 13624–13629, doi:10.1073/pnas.1110633108 (2011).

32 Li, Z. et al. Single-Copy Genes as Molecular Markers for Phylogenomic Studies in Seed Plants. Genome Biol Evol 9, 1130–1147, doi:10.1093/gbe/evx070 (2017).

33 Smith, S. A., Moore, M. J., Brown, J. W. & Yang, Y. Analysis of phylogenomic datasets reveals conflict, concordance, and gene duplications with examples from animals and plants. BMC Evol Biol 15, 150, doi:10.1186/s12862-015-0423-0 (2015).

34 Delsuc, F., Brinkmann, H. & Philippe, H. Phylogenomics and the reconstruction of the tree of life. Nat Rev Genet 6, 361–375, doi:10.1038/nrg1603 (2005).

35 Maddison, W. P. Gene trees in species trees. Systematic Biology 46, 523–536 (1997).

36 Michel, R. & Smirnov, A. V. The genus Flamella Schaeffer, 1926 (Lobosea, Gymnamoebia), with description of two new species. Eur J Protistol, 400–410 (1999).

37 Dykova, I., Lom, J., Dvorakova, H., Peckova, H. & Fiala, I. Didymium-like myxogastrids (class Mycetozoa) as endocommensals of sea urchins (Sphaerechinus granularis). Folia Parasitol (Praha) 54, 1–12 (2007).

38 Fiore-Donno, A. M., Tice, A. K. & Brown, M. W. A Non-Flagellated Member of the Myxogastria and Expansion of the Echinosteliida. J Eukaryot Microbiol 66, 538–544, doi:10.1111/jeu.12694 (2019).

39 Kudryavtsev, A., Pawlowski, J. Squamamoeba japonican. g. n. sp. (Amoebozoa): a deep-sea amoeba from the Sea ofJapan with a novel cell coat structure. Protist 164, 13–23 (2013).

40 Goodkov, A. V. & Seravin, L. N. Ultrastructure of the ‘giant amoeba’ Pelomyxa palustris. III. The vacuolar system; its nature, organization, dynamics and functional significance. Tsitologiya 33, 17–25 (in Russian with English summary) (1991).

41 Leadbeater BCS & Green, J. The flagellates. (Taylor and Francis, 2000).

42 Mitchell, D. R. The evolution of eukaryotic cilia and flagella as motile and sensory organelles. Adv Exp Med Biol 607, 130–140, doi:10.1007/978-0-387-74021-8_11 (2007).

43 Cavalier-Smith, T. Early evolution of eukaryote feeding modes, cell structural diversity, and classification of the protozoan phyla Loukozoa, Sulcozoa, and Choanozoa. Eur J Protistol 49, 115–178, doi:10.1016/j.ejop.2012.06.001 (2013).

44 Cavalier-Smith, T. in The Flagellates. (eds S. Leadbeater & J. Green) (Taylor and Francis, 2000).

45 Stechmann, A. & Cavalier-Smith, T. Rooting the eukaryote tree by using a derived gene fusion. Science 297, 89–91 (2002).

46 Cavalier-Smith, T. Only six kingdoms of life. Proceedings of the Royal Society of London Series B-Biological Sciences 271, 1251–1262 (2004).

47 Cavalier-Smith, T. Protist phylogeny and the high-level classification of Protozoa. European Journal of Protistology 39, 338–348 (2003).

48 Heiss, A. A., Walker, G. & Simpson, A. G. The flagellar apparatus of Breviata anathema, a eukaryote without a clear supergroup affinity. Eur J Protistol 49, 354–372, doi:10.1016/j.ejop.2013.01.001 (2013).

49 Roger, A. J. & Simpson, A. G. B. Evolution: Revisiting the Root of the Eukaryote Tree. Current Biology 19, R165–R167 (2009).

50 Chistiakova, L. V., Miteva, O. A., Frolov, A. O. & Skarlato, S. O. [Comparative morphology of the subphilum Conosa Cavalier-Smith 1998]. Tsitologiia 55, 778–787 (2013).

51 Derelle, R. et al. Bacterial proteins pinpoint a single eukaryotic root. Proc Natl AcadSci U S A 112, E693–699, doi:10.1073/pnas.1420657112 (2015).

52 Spiegel, F. W. in Encyclopedia of Evolutionary Biology (ed Richard M. Kliman) 325–332 (Academic Press, 2016).

53 Fiz-Palacios, O. et al. Did terrestrial diversification of amoebas (amoebozoa) occur in synchrony with land plants? PLoS One 8, e74374, doi:10.1371/journal.pone.0074374 (2013).

54 Schuster, J. J. & Markx, G. H. Biofilm Architecture. Advances in Biochemical Engineering/Biotechnology, 77–96, doi:doi:10.1007/10_2013_248 (2013).

55 Rich, V. I. & Maier, R. M. Aquatic Environments. Third Edition edn, (2015).

56 Allwood, A. C., Walter, M. R., Kamber, B. S., Marshall, C. P. & Burch, I. W. Stromatolite reef from the Early Archaean era of Australia. Nature 441, 714–718, doi:10.1038/nature04764 (2006).

57 Noffke, N., Christian, D., Wacey, D. & Hazen, R. M. Microbially induced sedimentary structures recording an ancient ecosystem in the ca. 3.48 billion-year-old Dresser Formation, Pilbara, Western Australia. Astrobiology 13, 1103–1124, doi:10.1089/ast.2013.1030 (2013).

58 Andersson, A. et al. Predators and nutrient availability favor protozoa-resisting bacteria in aquatic systems. Sci Rep 8, 8415, doi:10.1038/s41598-018-26422-4 (2018).

59 Matz, C. & Kjelleberg, S. Off the hook--how bacteria survive protozoan grazing. Trends Microbiol 13, 302–307, doi:10.1016/j.tim.2005.05.009 (2005).

60 Andersen, P. & Fenchel, T. Bacterivory by microheterotrophic flagellates in seawater samples. Limnol. Oceanogr. 30, 198–202. (1985).

61 Fenchel, T. The Ecology of Heterotrophic Microflagellates., Vol. 9 (1986).

62 Purcell, E. M. Life at low Reynolds number. American Journal of Physics 45 (1977).

63 Fenchel, T. Ecology of Protozoa. (Springer-Verlag, 1987).

64 Butler, H. & Rogerson, A. Consumption rates of six species of marine benthic naked amoebae (Gymnamoebia) from sediments in the Clyde Sea area. Journal of the Marine Biological Association of the United Kingdom 77, 989–997. (1997).

65 Jackson, S. M. & Jones, E. B. G. Interactions within biofilms: the disruption of biofilm structure by protozoa. Kieler Meeresforsch. Sonderh. 8, 264–268. (1991).

66 Martin, K. H., Borlee, G. I., Wheat, W. H., Jackson, M. & Borlee, B. R. Busting biofilms: free-living amoebae disrupt preformed methicillin-resistant Staphylococcus aureus (MRSA) and Mycobacterium bovis biofilms. Microbiology 166, 695. (2020).

67 Jahnke, J., Wehren, T. & Priefer, U. B. vitro studies of the impact of the naked soil amoeba Thecamoeba similis Greef, feeding on phototrophic soil biofilms. In Eur J Soil Biol 43, 14–22 (2007).

68 Anderson, O. R. Naked amoebae in biofilms collected from a temperate freshwater pond. J Eukaryot Microbiol 60, 429–431, doi:10.1111/jeu.12042 (2013).

69 Rogerson, A., Anderson, O. R. & Vogel, C. re planktonic naked amoebae predominately floc associated or free in the water column? Journal of Plankton Research 25, 1359–1365 (2003).

70 Parry, J. D. Protozoan grazing of freshwater biofilms. Adv Appl Microbiol 54, 167–196, doi:10.1016/S0065-2164(04)54007-8 (2004).

71 Wang, Y., Coleman-Derr, D., Chen, G. & Gu, Y. Q. OrthoVenn: a web server for genome wide comparison and annotation of orthologous clusters across multiple species. Nucleic Acids Res 43, W78–84, doi:10.1093/nar/gkv487 (2015).

72 Katoh, K. & Standley, D. M. MAFFT multiple sequence alignment software version 7: improvements in performance and usability. Mol Biol Evol 30, 772–780, doi:10.1093/molbev/mst010 (2013).

73 Capella-Gutierrez, S., Silla-Martinez, J. M. & Gabaldon, T. trimAl: a tool for automated alignment trimming in large-scale phylogenetic analyses. Bioinformatics 25, 1972–1973, doi:10.1093/bioinformatics/btp348 (2009).

74 Nguyen, L. T., Schmidt, H. A., von Haeseler, A. & Minh, B. Q. IQ-TREE: a fast and effective stochastic algorithm for estimating maximum-likelihood phylogenies. Mol Biol Evol 32, 268–274, doi:10.1093/molbev/msu300 (2015).

75 Edgar, R. C. Search and clustering orders of magnitude faster than BLAST. Bioinformatics 26, 2460–2461, doi:btq461 [pii] 10.1093/bioinformatics/btq461 (2010).

76 Chen, F., Mackey, A. J., Stoeckert, C. J. J. & Roos, D. S. OrthoMCL-DB: querying a comprehensive multi-species collection of ortholog groups. Nucleic Acids Res. 1, 34, doi:doi: 10.1093/nar/gkj123. (2006).

77 Stamatakis, A., Ludwig, T. & Meier, H. RAxML-III: a fast program for maximum likelihood-based inference of large phylogenetic trees. Bioinformatics 21, 456–463 (2005).

78 Zhou, X. et al. Quartet-Based Computations of Internode Certainty Provide Robust Measures of Phylogenetic Incongruence. Systematic Biology 69, 308–324 (2020).

79 Kobert, K., Salichos, L., Rokas, A., Stamatakis, A. &, C. t. I. C. a. R. M. f. P. G. T. Computing the Internode Certainty and Related Measures from Partial Gene Trees. Molecular Biology and Evolution 33, 1606–1617, doi:https://doi.org/10.1093/molbev/msw040 (2016).

80 Shimodaira, H. An approximately unbiased test of phylogenetic tree selection. Systematic Biology 51, 492–508 (2002).

81 Kishino, H., Miyata, T. & Hasegawa, M. Maximum likelihood inference of protein phylogeny and the origin of chloroplasts. (). https://doi.org/10.1007/BF02109483. J Mol Evol 31, 151–160 (1990).

